# Functional archaic DNA regulates molecular variation and is associated with disease risk across global populations

**DOI:** 10.1101/2023.04.26.538367

**Authors:** Jianning Kang, Aimee S Ramgolam, Louise Le Vot, Robert S. Young

## Abstract

The human genome contains many remnants of its evolutionary history, including a large number of evolutionarily volatile loci which have been introduced since our divergence from primates. One particularly intriguing source of novel DNA sequences is introgression events with archaic species which co-existed with modern humans. Both Neanderthals, who were common in Europe, and Denisovans, who have been observed only in Asia, have contributed genetic variants to the modern human genome but the functional consequences of these introgressed variants have yet to be investigated systematically. In this work, we show that Neanderthal and Denisovan DNA is most enriched for genetic variants which regulate gene expression in Europe and East Asia respectively, i.e. the populations in which the introgression event(s) most contributed to contemporary genetic variation. Neanderthal eQTLs, in particular, frequently upregulate gene expression. Archaic eQTLs from these two species regulate target genes with similar molecular functions which are distinct in each contemporary population, with the only common enrichment being for Neanderthal eQTLs to regulate taste receptor genes in both Europe and East Asia. We observed a correlated pattern of enrichment and depletion of medical phenotypes across Neanderthal and Denisovan eQTLs, including a shared enrichment for CNVs associated with developmental delay. Our results demonstrate the role of functional archaic DNA in regulating molecular phenotypes and disease risk across global populations and confirm the relevance of recently acquired DNA to contemporary human genetic variation.

**Author Summary:** Modern humans co-existed and interbred with two archaic human species (Neanderthals and Denisovans). The results of these events can still be detected as introgressed, archaic DNA sequences within the modern human genome. Here, we surveyed the contribution of functional archaic DNA across European and Asian populations by assessing their contribution to genetic variants which regulate gene expression in these two populations. We found that both species make a disproportionate functional contribution to the population with which they shared the most overlap (i.e. Neanderthals in Europe and Denisovans in East Asia). Although only Neanderthal DNA drives a higher level of gene expression compared to modern genetic variants, the DNA from both archaic species frequently regulates genes involved in many different biological processes and risk of disease, including a shared contribution to developmental delay. These results confirm the relevance of our recent evolutionary past in generating functional variation across global populations and the importance these recently introduced genetic sequences play in regulating current biological variation, such as disease risk.

## Introduction

Biological diversity across individuals, populations and species is thought to be primarily driven not by variation in their protein-coding gene complement, but in how the expression of these genes is regulated [1]. This view is supported by the observation that the overwhelming majority of genetic variants associated with phenotype through genome-wide association studies (GWAS) are found outside the borders of the protein-coding genes [2] and are enriched within transcriptional regulatory elements such as promoters and enhancers [3,4]. Many of these regulatory loci are evolutionarily volatile [5]. For example, it has been estimated that over 50% of orthologous genes shared between human and mouse have experienced the evolutionary turnover of at least one functional promoter since the divergence of their last common ancestor [6] approximately 75 million years ago [7]. Those promoters which have been gained along the human lineage arise from the insertion of transposable elements, are generally found in a repressive chromatin environment [8] and are enriched for expression QTLs (eQTLs, genetic variants which regulate gene expression) [9], confirming their importance in regulating human biology.

More recently, the human genome has also received evolutionary additions from archaic hominid species through introgression events during the time these species co-existed. Neanderthals were an archaic species of human who lived in Eurasia and were characterised by a robust build and squat stature compared to their modern-day counterparts [10]. They lived from approximately 130,000 [11] to 40,000 years ago [12], although the exact cause of their demise remains controversial. Multiple introgression events took place between Neanderthal and anatomically modern human (AMH) individuals which has resulted in ∼2% of the contemporary Eurasian genome being derived from that of the Neanderthal [13]. It has been suggested that Neanderthals were well adapted to the European environment and that this introgression may have therefore introduced beneficial alleles to recently arrived anatomically modern humans (AMHs) in a process known as ‘adaptive introgression’. This Neanderthal DNA has contributed to variation in a variety of traits including skin and hair colour, height and several behavioural traits [14]. Nevertheless, the Neanderthal population was also thought to be relatively small and suffered from inbreeding depression [15] which allowed the persistence of weakly deleterious alleles. This is exemplified by the presence of a Neanderthal-inherited locus on chromosome 3 which was identified as a major genetic risk factor for severe COVID-19 [16].

A second archaic species, the Denisovans, has been identified more recently. Only limited Denisovan remains have been identified, where they are thought to have lived from Siberia to Southeast Asia [17]. They lived until at least 30,000 years ago [18]. There is thought to have been two pulses of introgression between Denisovan and AMH individuals [19]. It is estimated that up to 6% of DNA in contemporary East Asian populations may have persisted from these introgression events [20].

The consequences of these introgression events for the modern human genome have been the focus of much investigation. Both Neanderthal and Denisovan DNA is largely found outside protein-coding regions but enriched within transcriptional regulatory sites including enhancers [21,22]. These observations suggest that, similar to sequences that have undergone changes throughout mammalian evolution, this introgressed archaic DNA is likely to have played a prominent role in rewiring our gene regulatory architecture. This has already been confirmed for Neanderthal genetic variants, which down-regulate gene expression across the brain [23]. Individual Neanderthal and Denisovan variants segregating within the Indonesian population have been demonstrated to contribute to local ancestry differences and differential gene expression across the Indonesian population [24].

While these studies have revealed the potential for introgressed archaic DNA to regulate gene expression, we wanted to investigate the scale of this across the genome and its potential to regulate phenotypes – both molecular and medical – across global populations. We performed a comprehensive study of the functional contribution of archaic DNA by integrating atlases of introgressed archaic alleles and expression quantitative trait loci (eQTLs). We next studied the contribution of these introgressed eQTLs to gene function and subsequent disease risk. Archaic eQTLs show clear signs of population stratification, where Neanderthal DNA is enriched for European eQTLs and Denisovan DNA for East Asian (Japanese) eQTLs. Those eQTLs which have persisted in contemporary human populations frequently have a large effect on gene expression across tissues. Archaic eQTLs regulate a variety of molecular functions. Archaic eQTLs display a range of enrichments and depletions for phenotype-associated genetic variants, although Denisovan eQTLs show a higher level of enrichment relative to Neanderthal eQTLs. The distribution of functional archaic DNA across populations therefore reflects human geographic history, where these variants regulate a wide variety of molecular and medical phenotypes, showing their continued importance to studies of human genetics.

## Results

### Neanderthal alleles are enriched over Denisovan at common variants in modern Europeans

We first investigated the distribution of archaic alleles in human populations in both Europe and East Asia. Variants obtained from Neanderthal and Denisovan samples were downloaded from publicly available databases (see Methods). The Denisovan genome has been sequenced to a much greater coverage (31-fold vs 1.3-fold for the Neanderthal Genome [25]) and this resulted in a much greater ability to detect genuine Denisovan variants. As in a recent study [22], we further required that these variants were not observed within the African population to remove variants that had been segregating prior to the emergence of archaic species such as Neanderthals and Denisovans. This resulted in a high-quality set of 13,191 and 442,873 Neanderthal and Denisovan variants, respectively, that we considered in subsequent analyses.

Consistent with their reported geographic distributions, we observed a clear enrichment of Neanderthal alleles relative to Denisovan in 1000 Genomes Project Europeans (Fig 1; odds ratio 1.2, Fisher’s exact test *p* = 1.4x10^-7^). We were able to recover this enrichment of Neanderthal alleles in Europe across a range of allele frequencies. This enrichment only became statistically non-significant above a minimum allele frequency of approximately 0.7 which is likely due to a reduced number of alleles and therefore reduced statistical power to detect deviations from neutrality, as shown by the large error bars. We also repeated this analysis but considered only alleles that were present in one of the two populations being considered here (see Methods). These population-specific alleles showed a comparable pattern to those for which we did not perform this filtering (S1 Fig).

**Fig 1.**
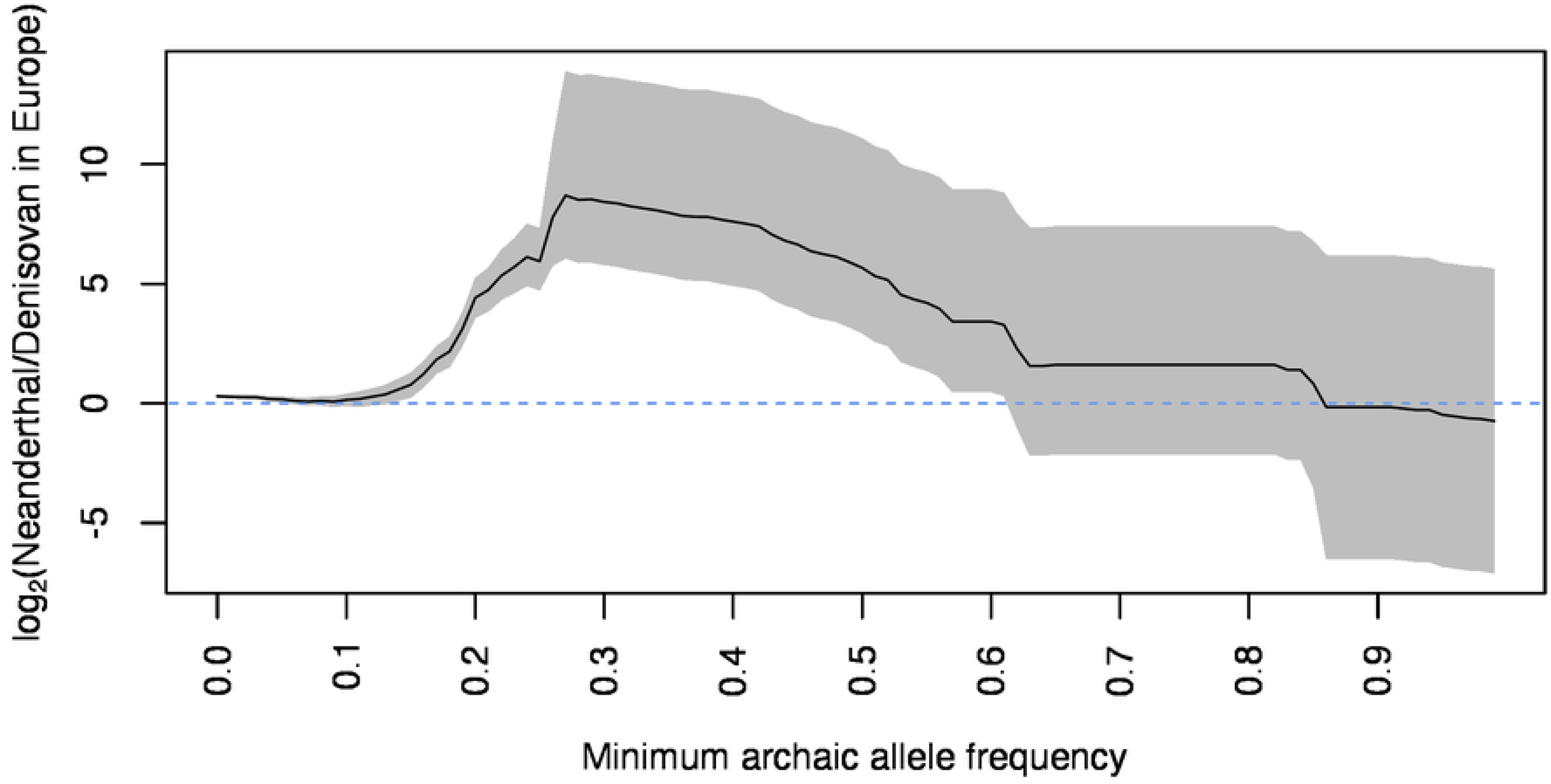
Neanderthal variants segregating within the European population are enriched over Denisovan variants. Odds ratio of the enrichment of Neanderthal over Denisovan variants segregating in the 1,000 genomes European super-population compared to the East Asian super-population calculated at increasing minimum archaic allele frequency. The grey curves indicate the 95% confidence interval of the odds ratio estimates. A log2(odds ratio) of 0 shown by the horizontal dashed blue line indicates an equal enrichment of Neanderthal alleles in Europe and East Asia.

It has been reported that Denisovan DNA is not present in the contemporary European population [22] which would appear to contradict these results. However, those alleles annotated here as being Denisovan in origin are segregating at a significantly reduced allele frequency in Europe (median ratio 0.6, Mann-Whitney *p* < 2.2x10^-16^). We speculate that these alleles are likely a result of migration and admixture between populations, introgressing the Denisovan variants from Asia to Europe across a number of generations, rather than suggesting that this provides evidence that Denisovan individuals were present in Europe. The continued presence of these variants across global populations nevertheless suggests that they play a functional role in regulating variation across individuals and are worthy of further investigation.

In order to balance maintaining a sufficient sample size with robust filtering of non-archaic derived sequences, we therefore considered all Neanderthal and Denisovan variants to be those which were observed in either Europe or East Asia, but not within the African population, to be genuine archaic variants which we considered in our subsequent analyses.

### Functional archaic DNA frequently upregulates gene expression across populations

Segregating alleles were defined as archaic if they overlapped either a Neanderthal or Denisovan variant, or modern if they did not overlap any sequenced Neanderthal or Denisovan nucleotide (see Methods). These locations were intersected with eQTL datasets from two distinct human populations to assess their functional impact. The GTEx eQTL atlas from the Genotype-Tissue Expression programme (GTEx) was first selected to have a strong contribution from the European population and these eQTLs were subsequently labelled as ‘European eQTLs’. We selected a Japanese eQTL atlas from immune cells [26] as a comparable set of East Asian eQTLs as it had the greatest coverage of samples that we are aware of within East Asian populations (n=6). The geographic history of each species has a clear effect on their contemporary eQTL contributions. When we considered variants segregating in Europe, Neanderthal DNA is significantly enriched only for European eQTLs while Denisovan DNA shows a greater enrichment for East Asian eQTLs (Fig 2). The same pattern was observed when considering Variants segregating in East Asia showed a similar pattern, where Denisovan variants were enriched for East Asian eQTLs but depleted for European eQTLs (S2 Fig, S1 Table) while Neanderthal eQTLs were similarly enriched in both datasets.

**Fig 2.**
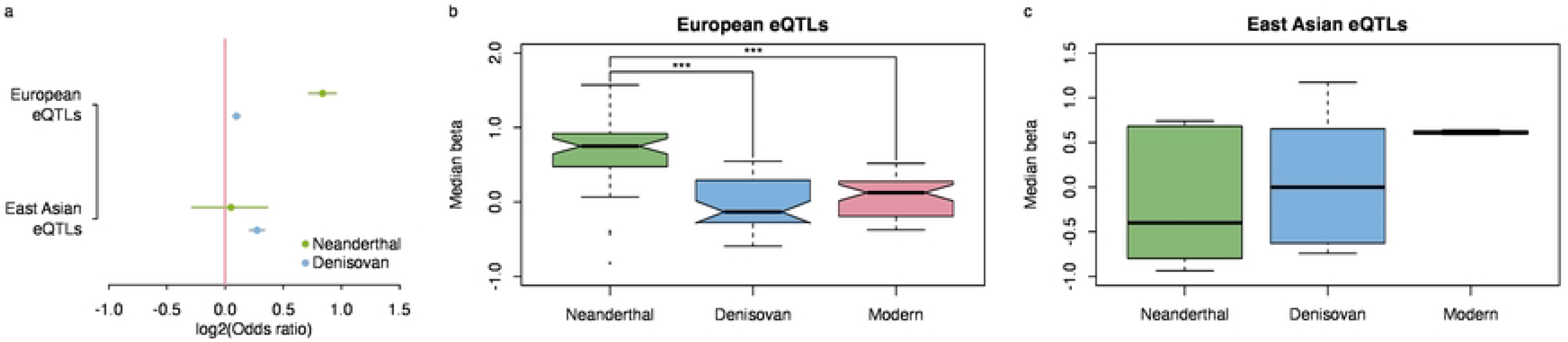
Neanderthal variants frequently up-regulate gene expression in Europe while Denisovan variants are enriched for East Asian eQTLs. (a) Odds ratio of the number of Neanderthal (in blue) and Denisovan (in blue) variants which harbour European and East Asian eQTLs relative to modern variants (purple line, x=0) when considering variants segregating within the contemporary European population. Circles indicate the point estimate of each odds ratio and horizontal lines indicate the 95% confidence interval of each estimate. (b and c) Distribution of median effect sizes across tissues (b, in Europe) and immune samples (c, in East Asia) for archaic and modern eQTLs. *** indicates Mann-Whitney test *p*-values < 0.001 when comparing median beta coefficients across tissues. All comparisons for East Asian eQTLs were nonsignificant (*p* > 0.05).

Similar patterns of relative Neanderthal enrichments in European eQTLs and Denisovan enrichments in East Asian eQTLs was noted across individual tissues and cell types (S3 and S4 Figs). While the East Asian dataset contained only immune cells and therefore was the only contribution to the Denisovan enrichments observed here, we were not able to detect any similar enrichments across any of the immune tissues in the European dataset using variants segregating either in Europe or East Asia (S3 and S4 Figs; S1 Table). None of the results were driven by the greater coverage of Denisovan DNA in the modern human genome, as similar patterns were obtained when down-sampling these variants to the same number as of Neanderthal variants (S5-8 Figs). The geographic history of these two species is therefore more important than tissue-specific factors in understanding this archaic contribution to contemporary human regulatory variation.

We next investigated the impact of archaic eQTLs on gene expression. The evolutionary history of each variant was reconstructed such that their reported effect sizes reflect the effect of the archaic or modern allele (see Methods). Across the European tissues, the Neanderthal eQTLs were found to have a greater, and more positive, effect on driving expression of their target genes relative to both modern eQTLs (Fig 2b, Mann-Whitney *p* = 9.3x10^-13^) and Denisovan eQTLs (Mann Whitney *p* = 7.0x10^-11^). Perhaps due in part to the smaller number of samples surveyed in the East Asian eQTL atlas (44 *vs* 6), we were not able to detect any significant differences between either set of archaic eQTLs and their modern counterparts in this dataset. The results presented here consider only alleles segregating in the European population but similar results were obtained when considering alleles segregating in East Asia (S4 Fig; S2 Table).

These results confirm that both archaic genomes have contributed a large number of regulatory variants which were disproportionately acquired or retained in the population which they had most geographic overlap with. Neanderthal variants which have persisted show a large effect size where they tend to increase the expression of their target genes.

### Archaic alleles regulate genes with a variety of molecular functions in both European and East Asian populations

We next investigated the characteristics of genes targeted by archaic eQTLs. In order to minimise sampling biases from genes regulated in individual tissues or cell lines, these analyses were performed by combining all eQTLs from either Europe or East Asia into a single dataset. Gene Ontology (GO) enrichment analysis revealed a number of significantly enriched terms within genes targeted by Neanderthal, Denisovan and modern eQTLs in both datasets (Fig 3, S9 Fig). Many terms showed similar enrichments for genes regulated by eQTLs from all three of these categories. The number of immune-related terms reported from the East Asian eQTL dataset likely reflects their sampling of immune cell types in contrast to the European dataset which covers a range of tissues.

**Fig 3.**
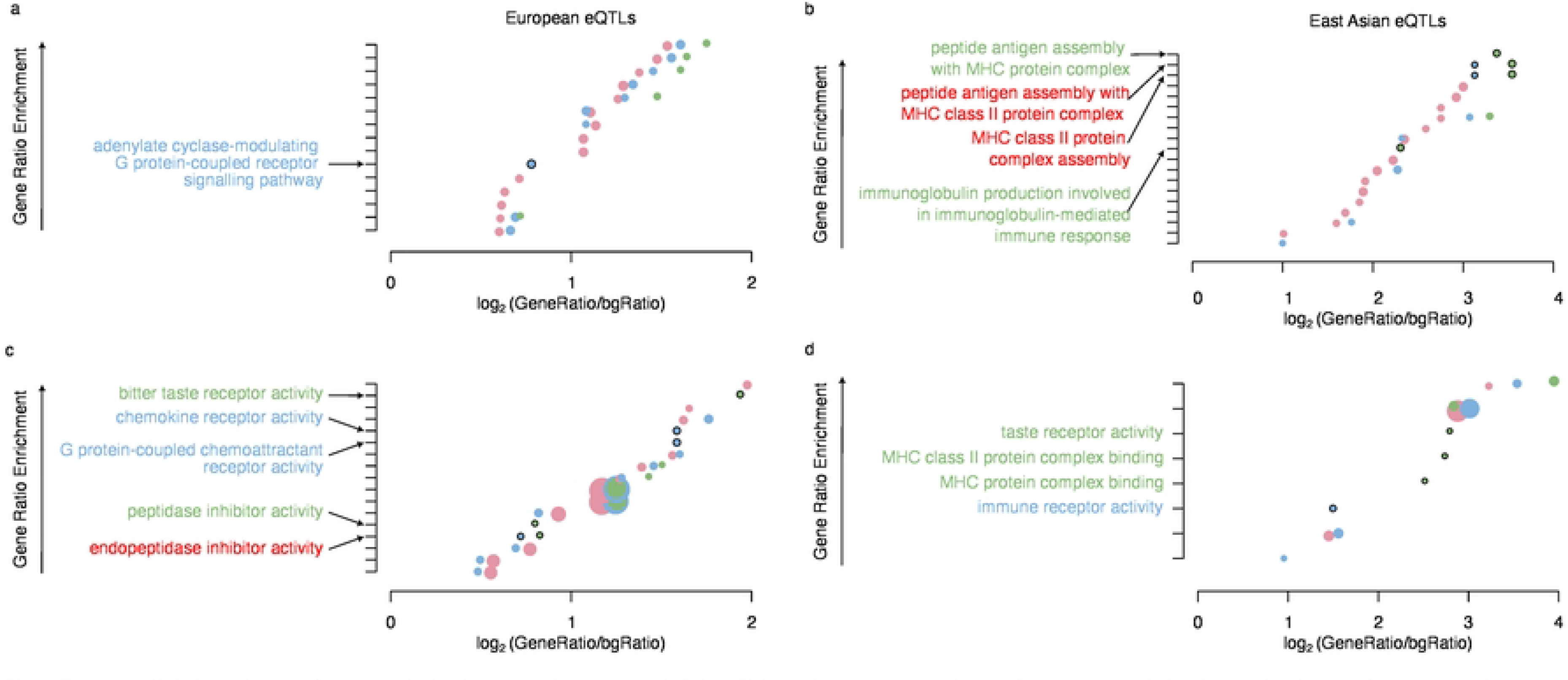
Archaic eQTLs regulate a variety of molecular functions. (a and b) Enrichment of Gene Ontology biological process terms within genes regulated by Neanderthal (green), Denisovan (blue) and modern (pink) eQTLs observed in Europe (a) and East Asia (b). Each circle shows a significant (Benjamini-Hochberg corrected *p*-value < 0.05) GO term where the size of the circle is proportional to the reported *p*-value. Solid lines around circles indicate those terms which are nominally enriched within archaic-regulated genes relative to their modern counterparts (Fisher’s exact test, *p* < 0.05). Only nominally-enriched terms are displayed on each y-axis, with those labels marked in red indicating terms that are nominally enriched for both Neanderthal and Denisovan eQTLs. (c and d) As above but for molecular function Gene Ontology terms.

We were unable to detect any terms which were significantly-differentially represented between the Neanderthal and Denisovan eQTLs. We did not attempt down-sampling of the Denisovan eQTLs as elsewhere in the study due to this lack of genome-wide significance. However, a number of the terms identified above (those shown in black-bordered circles) were nominally significantly enriched (Fisher’s exact test *p* < 0.05) relative to those genes which are regulated only by modern eQTLs. These enriched terms were largely distinct in the two contemporary populations, e.g. Neanderthal and Denisovan eQTLs in Europe were enriched for regulating endopeptidase inhibitor activity while both species were enriched in the East Asian eQTLs for MHC class II protein complex assembly. Only one term related to taste receptors could be detected as being significantly enriched for Neanderthal eQTLs in both Europe and East Asia (Fig 3c, d).

### Denisovan eQTLs make a greater contribution than Neanderthal eQTLs to disease risk

Given the importance of archaic eQTLs in gene regulation, we reasoned that they might play a similar role in regulating genes which were associated with medically important phenotypes. As in our previous work [9], we interrogated the relationship between archaic eQTLs and phenotype-associated genetic variants reported by association and family-based studies. We observed a wide range of enrichments and depletions when comparing the distribution of archaic to modern eQTLs across this diverse dataset of annotated variant collections (Fig 4; S3 Table). The distribution of Neanderthal and Denisovan eQTLs showed a weak correlation (Spearman’s rho 0.5, *p* = 3.1x10^-2^) implying that these two datasets might regulate common biological processes, consistent with their similar GO enrichments (Fig 3). Interestingly, the Denisovan eQTLs showed a consistently greater enrichment in contrast with their relative depletion of eQTLs relative to the Neanderthal contribution (Fig 2). We confirmed that this Denisovan enrichment was not down to a greater power to detect associations within Denisovan eQTLs by down-sampling this to the same number as Neanderthal eQTLs, where we continued to observe a significant correlation in the same direction (S10 Fig, Spearman’s rho < 0.7, *p* < 3.1x10^-7^).

**Fig 4.**
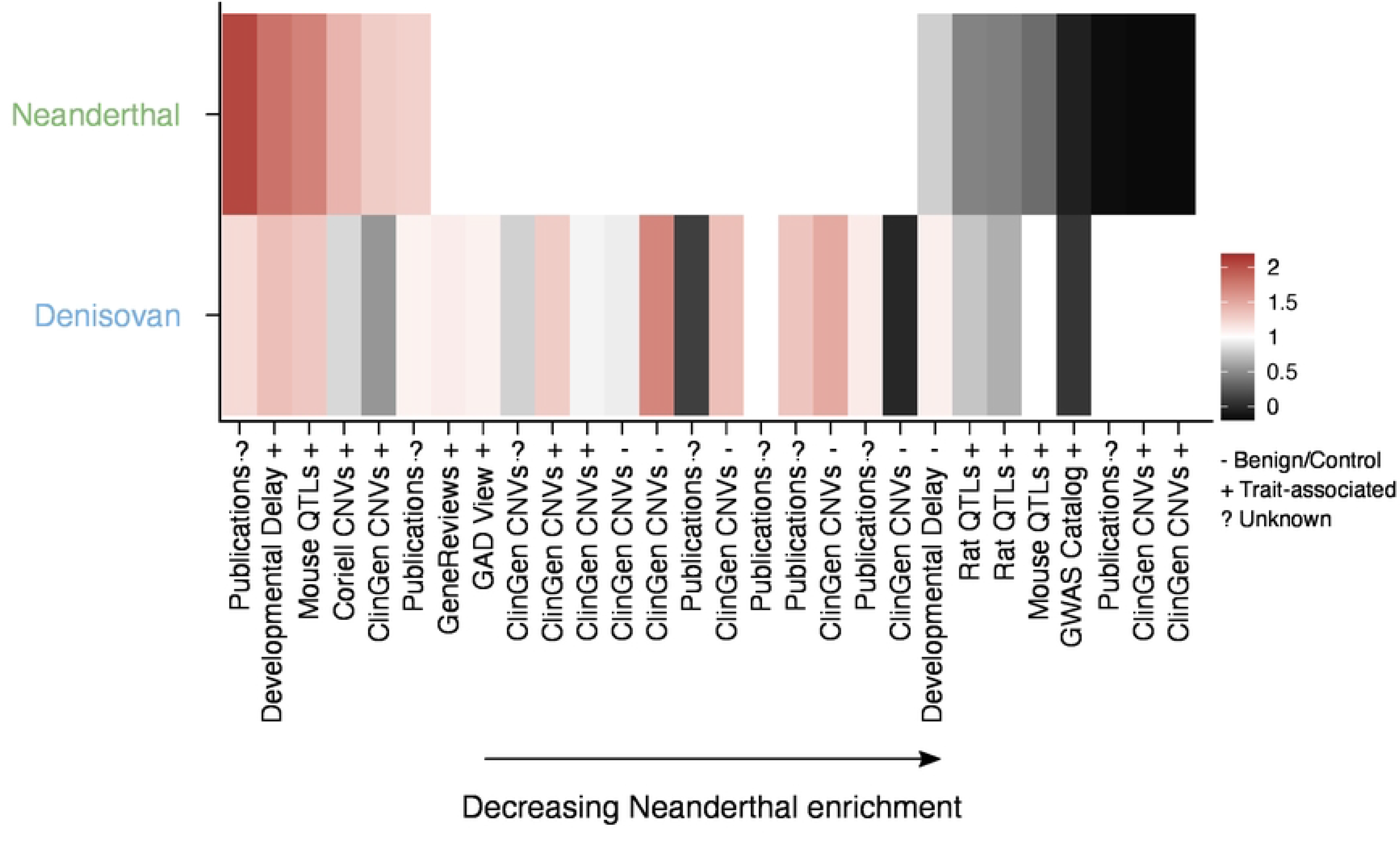
Enrichment of archaic eQTLs relative to their modern counterparts across phenotype- associated variant collections. Genetic variant collections (x-axis) were extracted from the ’Phenotype and Literature’ dataset group from the UCSC Genome Browser and are marked as ’benign’, ’trait- associated’ or ’unknown’ using each table description. Odds ratios were calculated using Fisher’s exact tests where red denotes eQTL enrichment relative to modern eQTLs and grey depletion. Non-significant (*p* > 0.05) associations have odds ratios rounded to one and are displayed as white.

We looked in more detail at the individual variant collections analysed here. Archaic eQTLs showed a consistent depletion within the GWAS Catalogue (odds ratio :<S 0.2, Fisher’s exact test *p* :<S 1.3x10^-7^; S3 Table) suggesting that the major contribution of these eQTLs is not mediated through their harbouring of genetic loci which contribute directly to disease. In contrast, archaic eQTLs were enriched for a number of copy-number variant (CNV) collections which have been associated with phenotypes, including developmental delay (S3 Table). It may be that these genomic regions which are susceptible to large-scale mutations are also those able to tolerate the continued presence of introgressed regulatory alleles, rather than a direction association between archaic eQTLs and disease.

Similar results were obtained when we considered only European or East Asian eQTLs (S11- S14 Figs). Although we noted no enrichments of Neanderthal DNA within East Asian eQTLs across these phenotype-associated variant collections, there were no significant deviations in the reported odds ratios across populations for either Neanderthal or Denisovan eQTLs (Mann-Whitney *p* > 0.05). Future large-scale studies of the population stratification of phenotype associations such as these analysed here could potentially be used to yield greater insights into the contribution of functional archaic variants to disease risk across global populations.

## Discussion

The analyses presented here has revealed the ongoing contribution of functional introgressed archaic DNA to the modern human genome. These variants regulate gene expression, molecular phenotypes and medically relevant traits in anatomically modern humans (AMHs) across contemporary European and East Asian populations. The distribution of archaic eQTLs across these populations reflects their geographic history, with Neanderthal eQTLs more frequent in Europe than East Asia and Denisovan eQTLs showing the inverse pattern (Fig. 2). Neanderthal variants make a disproportionate contribution to driving increased gene expression while both species frequently regulate a variety of molecular phenotypes (Fig. 3) and are commonly associated with disease risk (Fig. 4).

Recent introgression events such as these may continue to play an important role in contemporary population differentiation and the identification of population-specific associations in modern human individuals.

While our results reflect the geographic history of both archaic species, we note that in both populations the introgressed DNA is more frequently functional (as displayed by the presence of an eQTL) which might reflect selective pressures to retain functional, adaptive DNA in the population where the archaic species had the most geographic overlap. Denisovan DNA has already been demonstrated to contribute to variation in local ancestry across Indonesia [24] and adaptation to high altitude within the Tibetan population [27]. Further population-wide genetic studies of global populations which have been previously underrepresented should reveal further examples where archaic DNA contributes to local population adaptation and stratification.

Our analyses have demonstrated the importance of archaic DNA in transcriptional regulation across a wide range of molecular phenotypes and tissues (S3-4 Figs). The only consistent association we found across populations was for Neanderthal eQTLs in both Europe and East Asia to regulate taste receptor genes. We also show that archaic eQTLs can overlap phenotype-associated genetic variants (Fig. 4) from various sources, suggesting that they may contribute to different disease risk, consistent with previous studies that considered all introgressed Neanderthal DNA [14,28]. Many of these phenotype-associated variants are CNVs and it remains to be established whether the archaic eQTLs harboured within these may carry the causal, regulatory variant for the phenotype under consideration. These findings should stimulate further research into novel biological roles of functional, introgressed archaic DNA and potential differences in the contribution of Neanderthal compared to Denisovan DNA.

Neanderthal eQTLs frequently lead to upregulation of gene expression (Fig. 2; S2 and S5-6 Figs) which is consistent with other recently acquired regulatory loci in the human genome. These sequences, which have been inserted during primate evolution, gradually lose their ability to drive transcription [8] suggesting that there may be an optimum moderate level of gene expression in the human genome. Whether Neanderthal eQTLs behave similarly or whether their regulatory potential remains more constant over time remains to be fully determined.

Future studies of human genetic variation, particularly those focused on noncoding regulatory loci, should include evolutionarily novel variants in their analyses. These results confirm the importance of recently acquired sequences that has been added to the human genome within the last 150,000 years and suggests that a lack of evolutionary conservation should not be taken as evidence for lack of biological function. While many protein-genes are deeply conserved across species, those noncoding regions which experience evolutionary changes may be those which more frequently regulate genetic and phenotypic variation [9]. The results presented here further suggest that noncoding loci may experience differential selective pressures to their protein-coding counterparts, which result in a different relationship between evolutionary conservation and phenotype associations across these separate regions.

Our results further highlight differences in the role of genetic variants which regulate molecular and medical phenotypes. As previously reported [29], the overlap between eQTLs and GWAS signals is relatively limited. The enrichments observed here for archaic variants confirm this, as Neanderthal DNA is enriched for European eQTLs (Fig. 2) while it is the Denisovan eQTLs that more frequently harbour phenotype-associated genetic variants, most of which have been defined in European-biased population studies (Fig. 4). Many GWAS signals are prioritised by their overlap with functional data, e.g. the presence of a matched eQTL location. However, this approach may miss many important genetic variants and the mechanisms by which they mediate their phenotypic consequences. Our results should encourage future GWAS and post-GWAS analyses to explore complementary functional genomics datasets beyond eQTL atlases to increase the proportion of GWAS signals which can be explained and for which mechanistic hypotheses can be generated.

Overall, our study has revealed the importance of functional archaic DNA in regulating a significant component of contemporary human biology. These results highlight the importance of recent evolutionary events in driving ongoing biologically and medically relevant genetic variation across populations.

## Materials and Methods

### Genetic variation

We obtained contemporary human genetic variation from the 1,000 genomes project at http://ftp.1000genomes.ebi.ac.uk/vol1/ftp/release/20130502/. Multi-allelic and structural (i.e. non-SNP) variants were removed. European and East Asian variants were defined as those with an allele frequency > 0 in the EUR and EAS super-populations, respectively. European variants with an allele frequency of 0 in the EAS super-population were defined as European-specific while East Asian variants with an allele frequency of 0 in the EUR super- population were defined as East Asian-specific. We used the ‘AA=’ flag to annotate the ancestral state of all genetic variants. Variants which did not contain an ancestral annotation were still considered when performing genomic intersections but were not included when annotating the evolutionary trajectories for modern and archaic eQTLs.

Neanderthal variants identified by the Neanderthal Genome Project were downloaded from ftp://ftp.ebi.ac.uk/pub/databases/ensembl/neandertal/CatalogOfChanges.tgz. Segregating SNPs stored with the combined_SNP_anno.ns.har.filter.tsv file were converted to bed format and the UCSC liftOver tool used to convert these genomic coordinates to the hg19 assembly for comparison with other datasets. Genuine-neanderthal introgressed variants were identified as those sites which were found at an allele frequency of 0 in the AFR 1,000 genomes super-population and where the neanderthal allele differed from the human allele. Only these positions were retained for subsequent analyses.

We obtained Denisovan variants directly from the Table Browser of the hg19 genome assembly from the UCSC Genome Browser. In order to maintain consistency with the Neanderthal variants, we retained only SNP variants and those which were found at an allele frequency of 0 in the AFR 1,000 genomes super-population. Subsequently, variants where neither allele for the Denisovan genotype (as recorded by the ‘GT=’ flag) could be identified were removed.

Modern alleles were defined as variants segregating in the 1,000 genomes project data that did not overlap any Neanderthal SNP (defined as all locations in combined_SNP_annon.ns.har.filter.tsv) or Denisovan variant downloaded from the Table Browser.

### Functional genetic variation: eQTLs

eQTLs from the GTEx consortium were obtained from the patched version 6 release. Significant SNP-gene pairs were downloaded from the GTEx portal (https://storage.googleapis.com/gtex_analysis_v6p/single_tissue_eqtl_data/GTEx_Analysis_v6p_eQTL.tar).

Japanese white blood cell eQTL datasets [26] were downloaded from https://humandbs.biosciencedbc.jp/en/hum0099-v1#hum0099.v1.eqtl.v1. Significant-SNP gene pairs were considered as those for which a q-value ≤ 0.05 was reported from the permutation tests stored within /eQTL_gene_level/permutation/ directory.

We only considered those eQTLs which had previously been reported by the 1,000 genomes project (regardless of which population they were found in) as genuinely segregating variants within the contemporary human population. Those eQTLs which did not overlap with a variant within the 1,000 genomes project were removed from the analyses.

### Functional genetic variation: Phenotype-associated variants

Phenotype-associated variants were accessed as in our previous work [9] as all tracks contained within the ‘Phenotype and Literature’ group downloaded from the UCSC Genome Browser. Variants from each individual track were merged into a single-unified set of intervals. All tracks are detailed within S4 Table.

### Gene Ontology

Variants were associated with target genes as annotated by Ensembl through each eQTL annotation. Only protein-coding genes were submitted for gene ontology (GO) enrichment. First, protein-coding Ensembl IDs were converted to their Entrez counterparts using the mapIDs function within the clusterProfiler R package. GO enrichment analyses was then performed on these Entrez IDs using the enrichGO function within the same R package using the Benjamini-Hochberg correction for multiple testing and an adjusted *p*-value cutoff of 0.05. We calculated the gene enrichment ratios as the fraction of genes in the foreground (e.g. genes regulated by a Neanderthal eQTL) which were annotated with a specific GO term relative to the fraction of genes in the background. For all analyses presented here, the background given was the universe of all possible genes in the human genome.

### Statistical analysis and data visualisation

Statistical tests were performed using the R statistical package (version 3.6.1). Mann-Whitney U tests were conducted using the wilcox.test function, Student’s t-test using the t.test function and Fisher’s exact test using the fisher.exact function. A pseudocount was added to each value when considering the overlap of archaic and modern eQTLs with phenotype-associated variant collections.

## Acknowledgements

We are grateful to Prof Jim Wilson, Usher Institute, University of Edinburgh for helpful comments throughout this work and critically reviewing an earlier draft of the manuscript.

## Author Contributions

Conceived and designed the experiments: RSY JK. Performed the experiments: JK ASR LLV. Analysed the data: RSY JK ASR LLV. Wrote the paper: RSY JK.

## Supporting Information

**S1 Fig. Neanderthal variants specifically segregating within the European population are enriched over Denisovan variants.** Odds ratio of the enrichment of Neanderthal over Denisovan variants segregating only in the 1,000 genomes European super-population compared to variants segregating only in the East Asian super-population calculated at increasing minimum archaic allele frequency. The grey curves indicate the 95% confidence interval of the odds ratio estimates. A log2(odds ratio) of 0 shown by the horizontal dashed blue line indicates an equal enrichment of Neanderthal alleles in Europe and East Asia.

**S2 Fig. Neanderthal variants frequently up-regulate gene expression in Europe while Denisovan variants are enriched for East Asian eQTLs.** (a) Odds ratio of the number of Neanderthal (in green) and Denisovan (in blue) variants which harbour European and East Asian eQTLs relative to modern variants (purple line, x=0) when considering variants segregating within the contemporary East Asian population. Circles indicate the point estimate of each odds ratio and horizontal lines indicate the 95% confidence interval of each estimate. (b and c) Distribution of median effect sizes across tissues (b, in Europe) and immune samples (c, in East Asia) for archaic and modern eQTLs. *** indicates Mann-Whitney test p-values < 0.001 when comparing median beta coefficients across tissues. All comparisons for East Asian eQTLs were nonsignificant (p > 0.05)

**S3 Fig. Neanderthal variants are frequently enriched for European eQTLs segregating in the European population relative to Denisovan variants across tissues.** Odds ratio of the number of Neanderthal (in yellow) and Denisovan (in blue) variants which harbour eQTLs relative to modern variants (purple line, x=0), rank ordered by increasing Neanderthal odds ratio separately in European and East Asian populations. Vertical lines indicate the point estimate of each odds ratio and horizontal lines indicate the 95% confidence interval of each estimate. The black boxes indicate the depletions when considering all eQTLs across tissues (Europe) and cell types (East Asia) together.

**S4 Fig. Neanderthal variants are frequently enriched for European eQTLs segregating in the East Asian population relative to Denisovan variants across tissues.** Odds ratio of the number of Neanderthal (in yellow) and Denisovan (in blue) variants which harbour eQTLs relative to modern variants (purple line, x=0), rank ordered by increasing Neanderthal odds ratio separately in European and East Asian populations. Vertical lines indicate the point estimate of each odds ratio and horizontal lines indicate the 95% confidence interval of each estimate. The black boxes indicate the depletions when considering all eQTLs across tissues (Europe) and cell types (East Asia) together.

**S5 Fig. Neanderthal variants frequently up-regulate gene expression in Europe while Denisovan variants are enriched for East Asian eQTLs.** (a) Odds ratio of the number of Neanderthal (in yellow) and Denisovan (in blue) variants which harbour European and East Asian eQTLs relative to modern variants (purple line, x=0) when considering variants segregating within the contemporary European population. Denisovan variants have been down-sampled to the number of Neanderthal variants segregating within the European population. Circles indicate the point estimate of each odds ratio and horizontal lines indicate the 95% confidence interval of each estimate. (b and c) Distribution of median effect sizes across tissues (b, in Europe) and immune samples (c, in East Asia) for archaic and modern eQTLs. *** indicates Mann-Whitney test p-values < 0.001 when comparing median beta coefficients across tissues. All comparisons for East Asian eQTLs were nonsignificant (p > 0.05).

**S6 Fig. Neanderthal variants frequently up-regulate gene expression in Europe while Denisovan variants are enriched for East Asian eQTLs.** (a) Odds ratio of the number of Neanderthal (in yellow) and Denisovan (in blue) variants which harbour European and East Asian eQTLs relative to modern variants (purple line, x=0) when considering variants segregating within the contemporary East Asian population. Denisovan variants have been down-sampled to the number of Neanderthal variants segregating within the East Asian population. Circles indicate the point estimate of each odds ratio and horizontal lines indicate the 95% confidence interval of each estimate. (b and c) Distribution of median effect sizes across tissues (b, in Europe) and immune samples (c, in East Asia) for archaic and modern eQTLs. *** indicates Mann-Whitney test p-values < 0.001 when comparing median beta coefficients across tissues. All comparisons for East Asian eQTLs were nonsignificant (p > 0.05)

**S7 Fig. Neanderthal variants are frequently enriched for European eQTLs segregating in the European population relative to Denisovan variants across tissues.** Odds ratio of the number of Neanderthal (in yellow) and Denisovan (in blue) variants which harbour eQTLs relative to modern variants (purple line, x=0), rank ordered by increasing Neanderthal odds ratio separately in European and East Asian populations. Denisovan variants have been down- sampled to the number of Neanderthal variants segregating within the European population. Vertical lines indicate the point estimate of each odds ratio and horizontal lines indicate the 95% confidence interval of each estimate. The black boxes indicate the odds ratios observed when considering all eQTLs across tissues (Europe) and cell types (East Asia) together.

**S8 Fig. Neanderthal variants are frequently enriched for European eQTLs segregating in the East Asian population relative to Denisovan variants across tissues.** Odds ratio of the number of Neanderthal (in yellow) and Denisovan (in blue) variants which harbour eQTLs relative to modern variants (purple line, x=0), rank ordered by increasing Neanderthal odds ratio separately in European and East Asian populations. Denisovan variants have been down- sampled to the number of Neanderthal variants segregating within the East Asian population. Vertical lines indicate the point estimate of each odds ratio and horizontal lines indicate the 95% confidence interval of each estimate. The black boxes indicate the depletions when considering all eQTLs across tissues (Europe) and cell types (East Asia) together.

**S9 Fig. Archaic eQTLs regulate a variety of molecular functions.** (a and b) Enrichment of Gene Ontology biological process terms within genes regulated by Neanderthal (green), Denisovan (blue) and modern (pink) eQTLs in Europe (a) and East Asia (b). Each circle shows a significant (Benjamini-Hochberg corrected p-value < 0.05) GO term where the size of the circle is proportional to the reported p-value. Solid lines around circles indicate those terms which are nominally enriched within archaic-regulated genes relative to their modern counterparts (Fisher’s exact test, p < 0.05). (c and d) As above but for molecular function Gene Ontology terms.

**S10 Fig. Enrichment of archaic eQTLs down-sampled to contain the same number of Neanderthal and Denisovan eQTLs relative to their modern counterparts across phenotype-associated variant collections.** Genetic variant collections (x-axis) were extracted from the ’Phenotype and Literature’ dataset group from the UCSC Genome Browser and are marked as ’benign’, ’trait- associated’ or ’unknown’ using each table description. Odds ratios were calculated using Fisher’s exact tests where red denotes eQTL enrichment relative to modern eQTLs and grey depletion. Non- significant (p > 0.05) associations have odds ratios rounded to one and are displayed as white.

**S11 Fig. Enrichment of archaic European eQTLs relative to their modern counterparts across phenotype-associated variant collections.** Genetic variant collections (x-axis) were extracted from the ’Phenotype and Literature’ dataset group from the UCSC Genome Browser and are marked as ’benign’, ’trait-associated’ or ’unknown’ using each table description. Odds ratios were calculated using Fisher’s exact tests where red denotes eQTL enrichment relative to modern eQTLs and grey depletion. Non-significant (p > 0.05) associations have odds ratios rounded to one and are displayed as white.

**S12 Fig. Enrichment of East Asian archaic eQTLs relative to their modern counterparts across phenotype-associated variant collections.** Genetic variant collections (x-axis) were extracted from the ’Phenotype and Literature’ dataset group from the UCSC Genome Browser and are marked as ’benign’, ’trait-associated’ or ’unknown’ using each table description. Odds ratios were calculated using Fisher’s exact tests where red denotes eQTL enrichment relative to modern eQTLs and grey depletion. Non-significant (p > 0.05) associations have odds ratios rounded to one and are displayed as white.

**S13 Fig. Enrichment of archaic European eQTLs which have been down-sampled to contain the same number of Neanderthal and Denisovan eQTLs relative to their modern counterparts across phenotype-associated variant collections.** Genetic variant collections (x-axis) were extracted from the ’Phenotype and Literature’ dataset group from the UCSC Genome Browser and are marked as ’benign’, ’trait-associated’ or ’unknown’ using each table description. Odds ratios were calculated using Fisher’s exact tests where red denotes eQTL enrichment relative to modern eQTLs and grey depletion. Non-significant (p > 0.05) associations have odds ratios rounded to one and are displayed as white.

**S14 Fig. Enrichment of archaic East Asian eQTLs which have been down-sampled to contain the same number of Neanderthal and Denisovan eQTLs relative to their modern counterparts across phenotype-associated variant collections.** Genetic variant collections (x- axis) were extracted from the ’Phenotype and Literature’ dataset group from the UCSC Genome Browser and are marked as ’benign’, ’trait-associated’ or ’unknown’ using each table description. Odds ratios were calculated using Fisher’s exact tests where red denotes eQTL enrichment relative to modern eQTLs and grey depletion. Non-significant (p > 0.05) associations have odds ratios rounded to one and are displayed as white.

**S1 Table. No. archaic and ancestral eQTLs across various tissues.**

**S2 Table. Median beta effect size of eQTLs across archaic and ancestral eQTLs for the European and East Asian eQTL datasets.**

**S3 Table. No. ancestral and archaic eQTLs overlapping each genetic variant collection.**

**S4 Table. Web links for genetic variants reported by the 1,000 genomes project and phenotype-associated variant collections collected from the UCSC Genome Browser.**

